# Queryfuse is a sensitive algorithm for detection of gene-specific fusions

**DOI:** 10.1101/2020.03.15.993089

**Authors:** Yuxiang Tan

**Affiliations:** Boston University, United States

## Abstract

Recurrent chromosomal translocations, known as fusions, play important roles in carcinogenesis. They can serve as valuable diagnostic and therapeutic targets. RNA-seq is an ideal platform for detecting transcribed fusions, and computational methods have been developed to identify fusion transcripts from RNA-seq data. However, some transciptome realignment procedures for these methods are unnecessary, making this task computationally expensive and time consuming. Therefore, we have developed QueryFuse, a novel hypothesis-based algorithm that identifies gene-specific fusion from pre-aligned RNA-seq data. It is designed to help biologists quickly find and/or computationally validate fusions of interest, together with visualization and detailed properties of supporting reads. By aligning reads to Query genes at the pre-processing step with a more sensitive, memory intensive local aligner, QueryFuse can reduce alignment time and improve detection sensitivity.

QueryFuse performed better or at comparable levels with two popular tools (deFuse and TopHatFusion) on both simulated and well-annotated cell-line datasets. Finally, using QueryFuse, we identified a novel fusion event with a potential therapeutic implication in clinical samples. Taken together, our results showed that QueryFuse is efficient and reliable for detecting gene-specific fusion events.

## INTRODUCTION

A gene fusion, also referred to as chromosomal translocation, denotes the event whereby two normally separated genes are joined together as a consequence of DNA double strand breakages (DSBs), followed by a DNA repair error^1,2^. A special type of normal gene fusion called V(D)J recombination is the key factor in the process of antibody generation in the human immune system^3^. However, gene fusions are also known to play important roles in tumorigenesis and progression in basically all tumor types^1,4,5^. It has been shown that, on the one hand, the decrease or eradication of a disease-associated chimera can be a successful treatment for tumor removals, while manipulated gene fusions can give rise to neoplastic disorders ^1^.

Depending on the location of the fusion breakpoint, an actionable gene fusion can either lead to the formation of a new chimeric protein by changing the fusion genes’ reading frame or stop codon, or dysregulate the expression of fusion genes by exchanging regulatory elements with this pair of fusion genes^1,6^. On the other hand, a silence gene fusion will not transcribe the DNA change to RNA level. For example, in chronic myelogenous leukemia (CML), *BCR-ABL1* fusion produces a chimeric protein that induces growth factor independence and leukemogenesis^6,7^. Inhibition of the *BCR-ABL1* enzyme by imatinib stops cancer cell growth and results in significant improvement of survival rate (from 30% to 89%) for CML patients^8,9^. For the *ETS*-*TMPRSS2* fusion in prostate cancer, the androgen-responsive regulatory elements of *TMPRSS2* replace the elements of the *ETS* gene and upregulate the expression of the *ETS* family member^6,10^.

Because gene fusions involve a DNA DSB and mis-repair^1,2,6^, whole genome sequencing (WGS) with deep coverage is the ideal way to find all gene fusions (both actionable or silent) in a sample. However, WGS’s high cost prevents it from broad adoption in clinical practices. Instead, RNA sequencing (RNA-Seq) is the most cost-effective approach to detect clinically actionable fusions.

### RNA-Seq Based Fusion Detection on Expressed Genes

Analogous to WGS’s targeting of the whole genome, RNA-Seq targets only the RNA present at a given moment^11^. Because only a small portion of the genome is transcribed to be functional at any given time, with the same number of sequencing reads, RNA-Seq can provide much greater depth of coverage (>50 times) than WGS^12^. Consequently, RNA-Seq is much more sensitive than WGS for detecting low-frequency variations. Moreover, RNA-Seq identifies gene fusion transcripts, which can show how the protein product will change and/or what regulation elements are maintained in the fusion.

Since 2010, at least 21 RNA-Seq based fusion-detection tools have been published (Table 1). Generally, RNA-Seq based fusion-detection tools can be categorized into two groups: 1) fragment based approaches and 2) pseudo-reference based approaches (Table 1). Fragment based approaches split reads into fragments and align fragments to a regular reference genome/transcriptome. Pseudo-reference based approaches generate a pseudo-reference by *de novo* assembly and use it together with the regular reference^13^. Although most of the RNA-Seq based fusion-detection tools are designed specifically for fusion-detection, the rest are RNA-Seq aligners with integrated fusion-detection models (Table 1). Several reviews from various resources have previously given evaluations of fusion detection tools; however no single tool performed the best in all test datasets^14,15^.

**Table 1.**
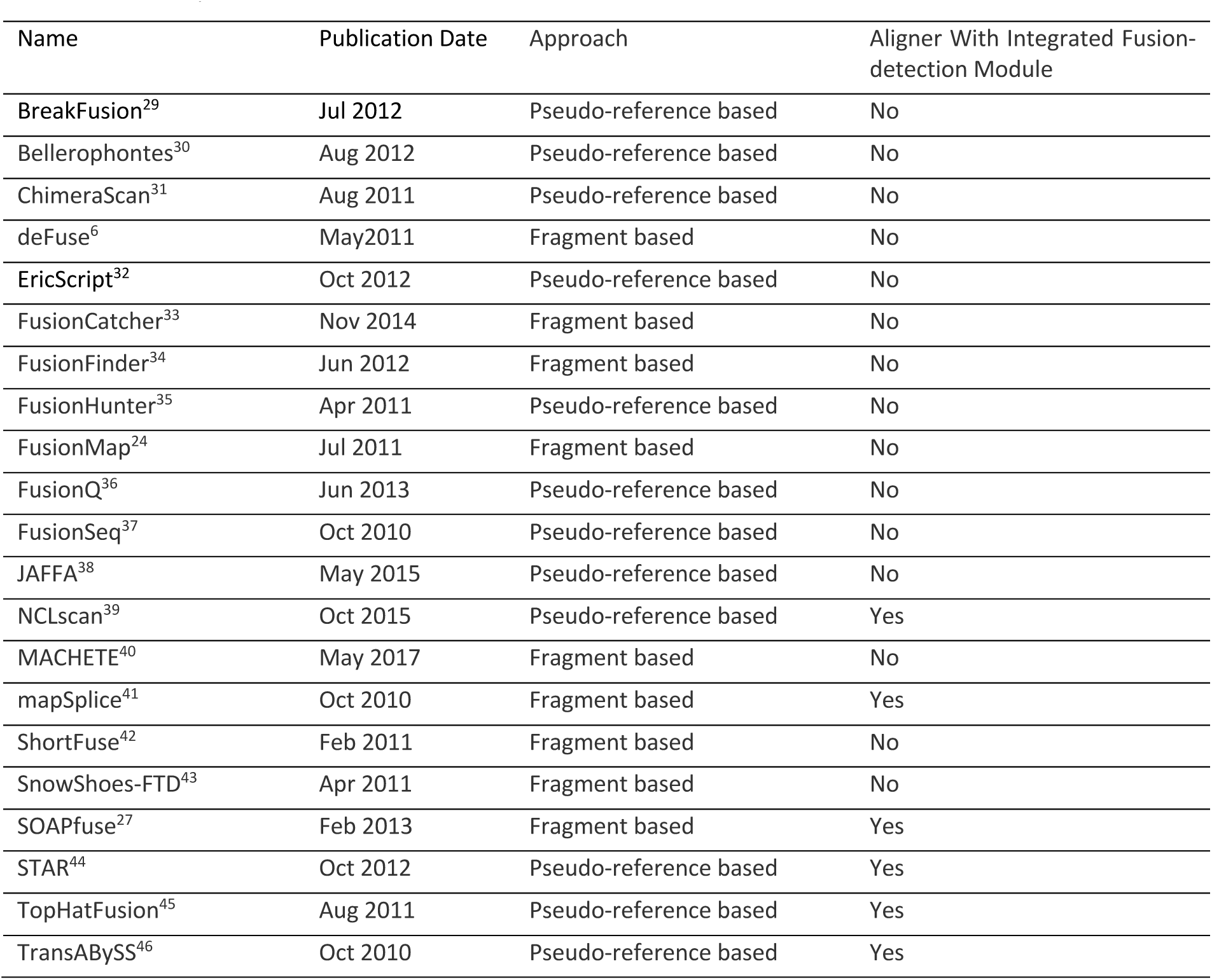
RNA-Seq based fusion-detection tools

Existing RNA-Seq based fusion-detection algorithms have two common features: 1) the use of raw reads as input (except BreakFusion), and 2) global fusion detection.

Regardless of downstream analysis (e.g., structure variation analysis, expression quantification analysis), alignment is consistently the first and most time-consuming step of RNA-Seq analyses, because it converts meaningless sequences into transcriptomic information. Consequently, because more than 90% of the reads are not fusion related in a regular RNA-Seq library, aligning all the raw reads for the purpose of fusion-detection only is inefficient and unnecessary. Therefore, fusion-detection tools could be more efficient by using already aligned data instead of using raw data.

It is true that global detection can reveal the big picture of all potential fusions. However, a long list of candidate events is overwhelming for most biologists. Since researchers have their own list of target genes/pathways, they will focus mostly on their selected lists, rather than on other unrelated candidates, especially when they have hypotheses generated by biological facts. Therefore, a fusion detection tool that focuses on only a selected list of genes will save researches’ time by eliminating “noise.”

### QueryFuse: A Gene Specific Fusion-Detection Algorithm

Global fusion-detection is generally time-consuming, has low recall and precision rates,^16^ and returns irrelevant fusions that act to distract biologists. With copy number variation (CNV) and gene expression data, biologists can now generate hypotheses regarding novel fusions with specific genes^1,5^, and this hypothesis driven fusion detection approach can serve as an alternative to global detection.

In this manuscript we present QueryFuse, a gene specific RNA-Seq based fusion-detection algorithm, designed to detect fusions of specific genes in a user-friendly, hypothesis focused, efficient, and reliable way for biologists. Unlike other fusion-detection algorithms (except BreakFusion), QueryFuse uses aligned BAM files as input in order to avoid the costly alignment step prevalent in other fusion detection tools. Moreover, instead of using a noisy global fusion detection method, QueryFuse focuses on the user’s gene list of interest, making it possible to utilize a memory intensive local aligner that improves recall and precision^16^. QueryFuse’s output includes details that allow users to better understand detected fusions, such as: the exact read ID of all fusion supporting reads, the alignment graph around the fusion breakpoint, and the values of ranking features. In the final report, all fusions are ranked by the sum of select features to help users prioritize their focus. In contrast to most fusion-detection algorithms, which use only the exon regions, QueryFuse covers all annotated regions (exons and introns) of genes.

Meanwhile, we found that many false positive (FP) fusions were discovered because the two breakpoints were too close to each other. To address this FP issue, we defined a gene fusion in QueryFuse as an event where two breakpoints are inter-chromosomal, or intra-chromosomal with more than 50 kb apart (the average gene density in the human genome is about 40-45 kb DNA per gene^12^).

## METHODS (900-1800: NOW 2300 WORDS, TOO MUCH?)

### The QueryFuse Algorithm

Essential terms used in this study are listed in Figure 1. A *fragment* is a contiguous sequence of nucleotides from a cDNA molecule. A *read* is a sequenced end of a fragment. We define *paired-end sequencing* as the protocol of sequencing both ends of the same fragment and the two sequenced ends as *paired ends*. The non-sequenced fragment portion between paired ends is called *insert region*, and its length *insert-size*. If the insert-size is negative, the paired ends overlap. We define as *paired-end-aligned (PEA)* those reads in which both members of the pair are properly aligned to the reference. Similarly, *single-end-aligned (SEA)* denotes read pairs where only one of the ends is properly aligned. A read is *unmapped* when it is not properly aligned. In paired-end sequencing, a SEA read should always have a corresponding unmapped read. It is also possible that both paired-ends are unmapped, which are called both-end-unmapped (BEU), especially when the insert-size is negative. We define *fusion boundaries* as the precise, nucleotide-level genomic coordinates of the breakpoints on both sides of the fusion gene pair.

**Figure 1.**
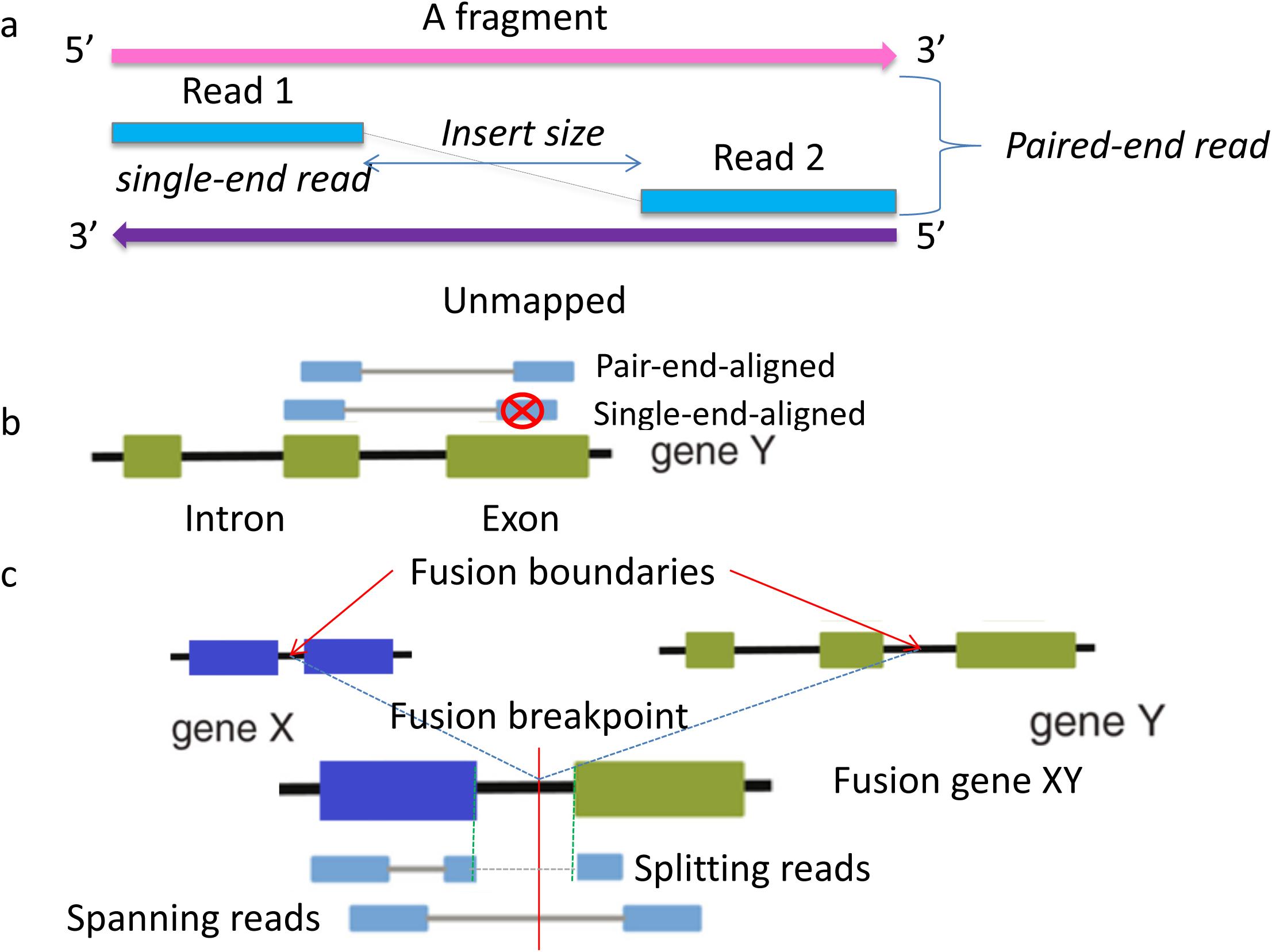
Definition of essential terms. a) A blue bar stands for a sequenced read. The pink and purple bars are two strands of a fragment. The arrow points in the sequencing direction, and a read is always on the 5’ end of a strand. If only one read is sequenced per fragment, it is single-end sequencing and the read is called ‘single-end read.’ If each fragment has two sequenced reads, it is paired-end sequencing and the pair of reads is called ‘paired-end reads.’ b) The green boxes stand for exons and the black strings stand for introns. For paired-end sequencing, if only one read of a pair is properly aligned to the reference (generally on exon in RNA-Seq data), it is called single-end-aligned (SEA). If both reads of a pair are properly aligned, it is called paired-end-aligned (PEA). c) The red string indicates a pair of fusion boundaries on gene X and gene Y (generally on intron in DNA). From RNA-Seq, a fusion gene XY will present as two exons fused together. S*panning* reads are those PEA reads that have fusion boundaries in the insert region; whereas, a *splitting* read denotes a read with the fusion boundaries within its length.

In fusion-supporting reads, there are two categories: spanning reads and splitting reads. Spanning reads are PEA reads that have fusion boundaries in their insert regions; whereas, *splitting* reads are reads with the fusion boundaries within their lengths.

### QueryFuse’s workflow

QueryFuse consists of three main steps (Figure 2). 1) Extraction of fusion candidate reads obtained from pre-aligned data using the query gene. By extracting fusion candidate reads from pre-aligned data using the query gene, we eliminated the previous step of realigning all the reads for fusion-detection. Therefore, QueryFuse saved considerable time and, to our knowledge, implemented existing information that was not previously used in other algorithms. 2) Classification of fusion candidate reads. Fusion candidate reads are carefully separated into a total of three different categories (Figure 3), with the reads in each category analyzed by category-specific methods aimed at maximizing fusion-detection precision and recall rates. 3) Result summarization. The results from all categories were merged. To increase accuracy, several filters were applied to remove false positives. To help user interpret the results, we prioritized reliable fusions by feature-specific ranking across detected fusion events. In the last step, summary reports and alignment graphs in text formats were automatically generated.

**Figure 2.**
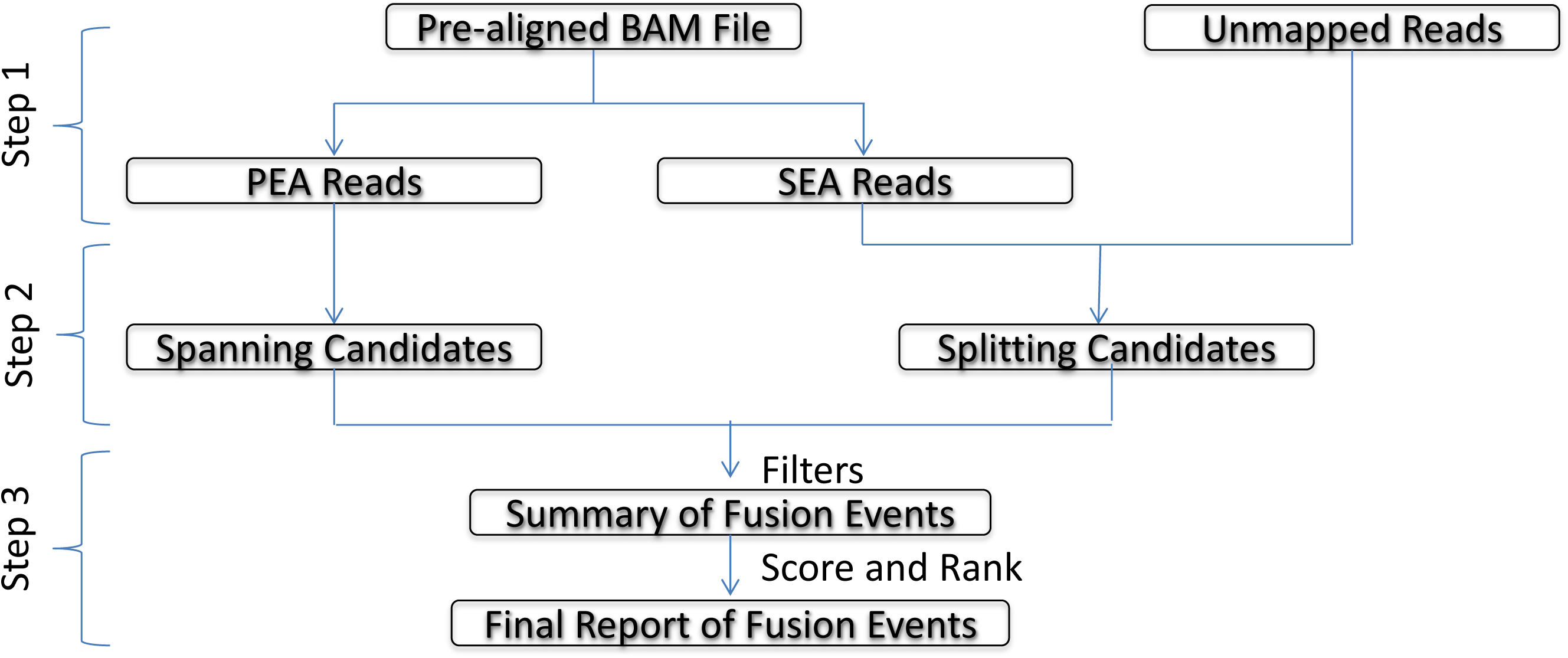
QueryFuse’s workflow. Step 1) Split the pre-aligned BAM file into PEA and SEA reads; Step 2) Extract spanning candidates from PEA reads and splitting candidates from SEA and unmapped reads; Step 3) These candidates are summarized and filtered to get the summary of fusion events, and output the final report of fusion events after scoring and ranking.

**Figure 3.**
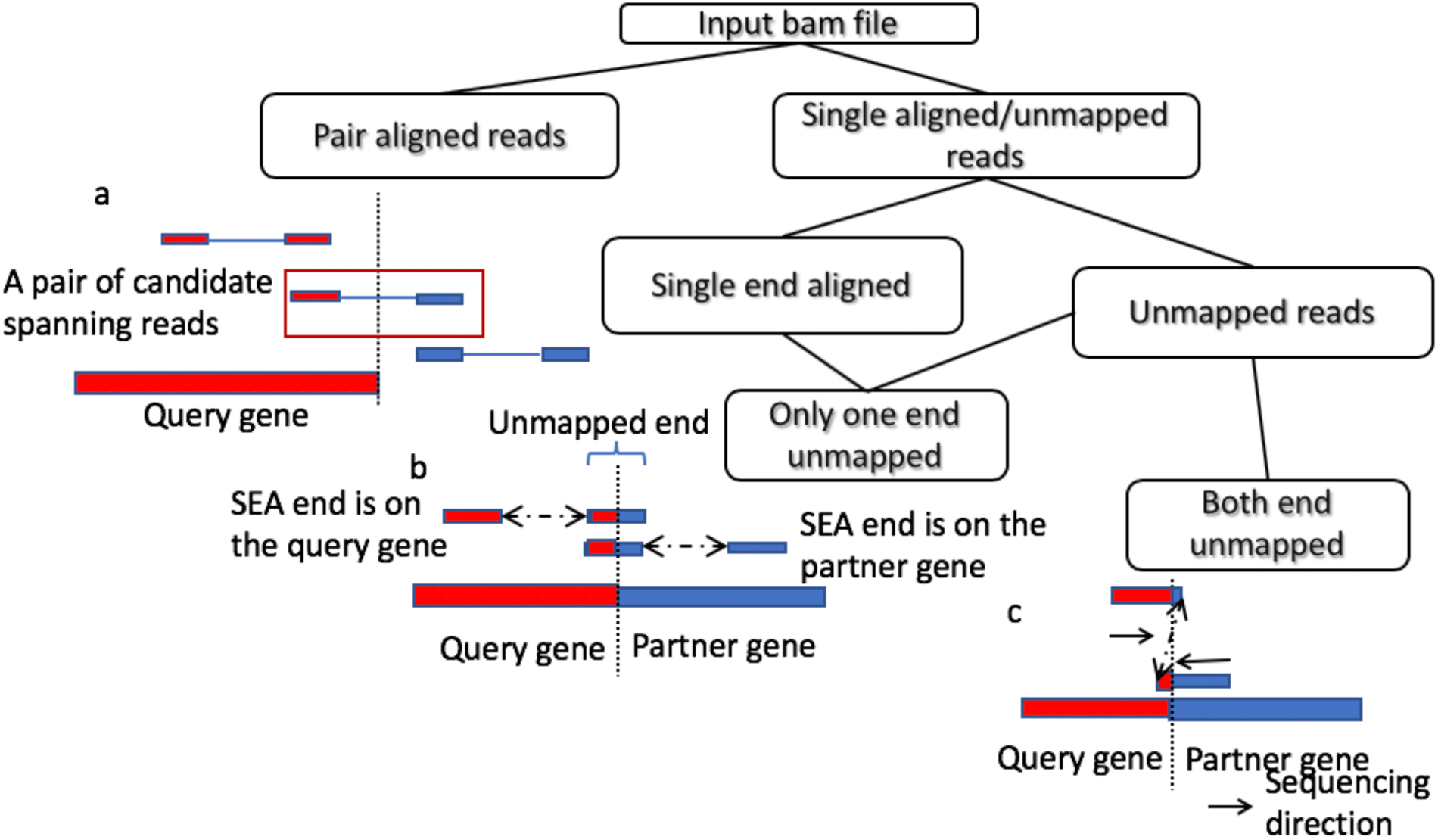
Categories of fusion candidate reads. Red boxes stand for the query gene and blue boxes stand for the partner gene (which is generally unknown at this stage). The dashed line stands for fusion boundaries. a) Paired-end processing category uses PEA reads as input and then extracts candidate spanning reads; these candidates have one end aligned to the query gene and the other end aligned to the partner gene(s); b) Single-end processing category uses SEA reads and their complementary unmapped reads as input. To be a pair of candidate splitting reads, the SEA reads can be on the query gene or on the partner gene(s), while their complementary unmapped reads must be partially on the query gene, with the other part on the partner gene(s); c) Unmapped pair processing category uses read pairs with both ends unmapped as input. To be a pair of candidate splitting reads, the two reads should overlap and fusion boundaries should be in the overlapped region. Additionally, the two reads must be sequenced from opposite directions. Meanwhile, if the major portion of one end of a pair is on the query gene, the major portion of the other end must be on the partner gene.

### Extraction of fusion candidate reads involving the query gene from pre-aligned data

QueryFuse was designed to detect all candidate fusion events involving a given gene of interest (the query gene) from pre-aligned RNA-Seq data. In this study, we used TopHat^17^-aligned data as our input. The aligned reads were separated into PEA and SEA reads. PEA reads partially aligned to the query gene were collected as spanning candidates. SEA and BEU reads partially aligned to the query gene from a local aligner and were extracted as splitting candidates. (Now we used BLAT^18^ as the local aligner, but other local aligners, such as BLAST^19^, and STELLAR^20^ could be supported in the future.)

### Classification of fusion candidates into different categories

Candidate fusion supporting reads were grouped into three categories (Figure 3).

The first category was paired-end processing (Figure 3a). Only the read pairs, which have one end aligned to the query gene and the other end aligned to other genes, are retained as spanning read pairs. The second category was single-end processing (Figure 3b), the most distinctive feature of QueryFuse algorithm. The key concept was: In a SEA read pair, gene information from the SEA end reliably located the breakpoint location on the unmapped end. Because query gene must be part of an unmapped end, the unmapped end should have two parts: ‘the piece from the query gene’ and ‘the piece from an unknown gene.’ Thus, we locally aligned the unmapped ends in the SEA read pairs to the query gene to obtain ‘the piece from the query gene.’ Depending on where the SEA end aligned, two subcategories (‘SEA end was on the query gene’ and ‘SEA end was on the partner genes) were used to confirm where ‘the piece from an unknown gene’ originated (Figure 3b). In the first subcategory, we globally aligned ‘the piece from an unknown gene’ to the genome; however, the result from this subcategory was less reliable than the second subcategory’s. To confirm that the unknown gene was the partner gene where the SEA end was detected in the second subcategory, we locally aligned ‘the piece from an unknown gene’ to the gene where the SEA end was detected.

The last category was unmapped pair processing (Figure 3c). In a read pair, if the two ends overlapped and the fusion breakpoint was in this overlapped region, neither end could be aligned. Since fusion events involving the query gene were those targeted, both ends of a read pair were locally aligned to the query gene. The unmapped portions were then globally aligned to the genome. When fusion boundaries were the same in a pair of genes, where both ends were aligned, this read pair was considered a splitting read pair. Additionally, we used the characteristics of sequencing direction and splitting lengths (Figure 3c) as criteria.

### Result Summarization

To summarize the results at event level, spanning and splitting reads for each fusion event were merged together by their alignment location. For each fusion event, since fusion boundaries of its splitting reads should be the same, QueryFuse (QF) used fusion boundaries of splitting reads (from the single-end processing category and unmapped pair processing category) to represent fusion events and keys to search for complementary spanning read pairs.

To evaluate the number of multiple-alignments for each fusion event, a consensus fusion sequence was generated using all supporting splitting reads (*N*) and was locally aligned to the genome. If the consensus sequence could be aligned equally well to *M* different locations in the genome, we considered these locations as multiple-alignments. The splitting reads number for each alignment location of this fusion event equaled to *N/M* (Figure 4). If the alignments on the query gene and fusion partner gene had an overlapped region, we defined this region as the ‘shifted region’ (which meant shifting the breakpoint location in this region would not change the fusion sequence) and its length as the ‘shifted range’ (Figure 5). The fusion breakpoint would be adjusted to the middle of this region. Otherwise, there was no fusion breakpoint adjustment for this fusion event because the ‘shifted range’ was zero. Additionally, the consensus sequence was essential to eliminate false positive, caused by sequence similarity. If the consensus sequence aligned well on the fusion partner gene (<23 bp unmapped), this fusion event was considered as a false positive. In this case, the consensus sequence could possibly only come from the fusion partner. It was falsely considered as a fusion candidate because the query gene had a long homologous region with this fusion partner. (A >30bp long sequence that occurs multiple times across the genome is considered a *high similarity* region.)

**Figure 4.**
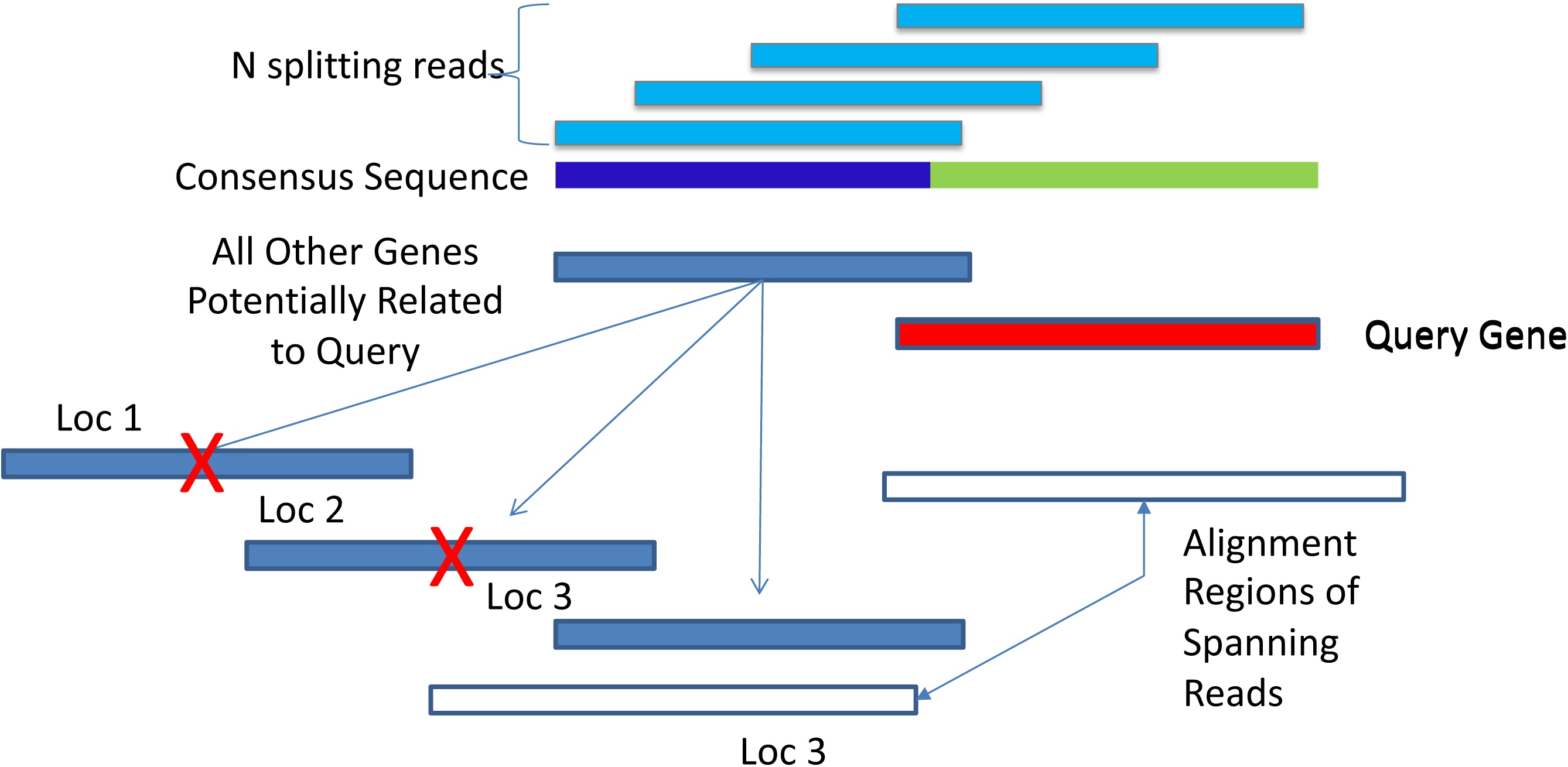
Multiple-alignment scenario. For each detected fusion event, all of its splitting reads (light blue) are used to build the representative consensus sequence of the fusion product from two genes. This consensus sequence must have a portion that can align to the query gene (red), while the other side of the consensus sequence aligns to other locations (dark blue) on the reference. If the number of location(s) is bigger than one, this fusion event has multiple-alignments. In this case, all locations (e.g. 3 in this case) are recorded and matched to alignment regions of corresponding spanning reads (white). Only the one(s) supported by alignment regions of most spanning reads (e.g. Loc 3) is extracted as the representative of the fusion event.

**Figure 5.**
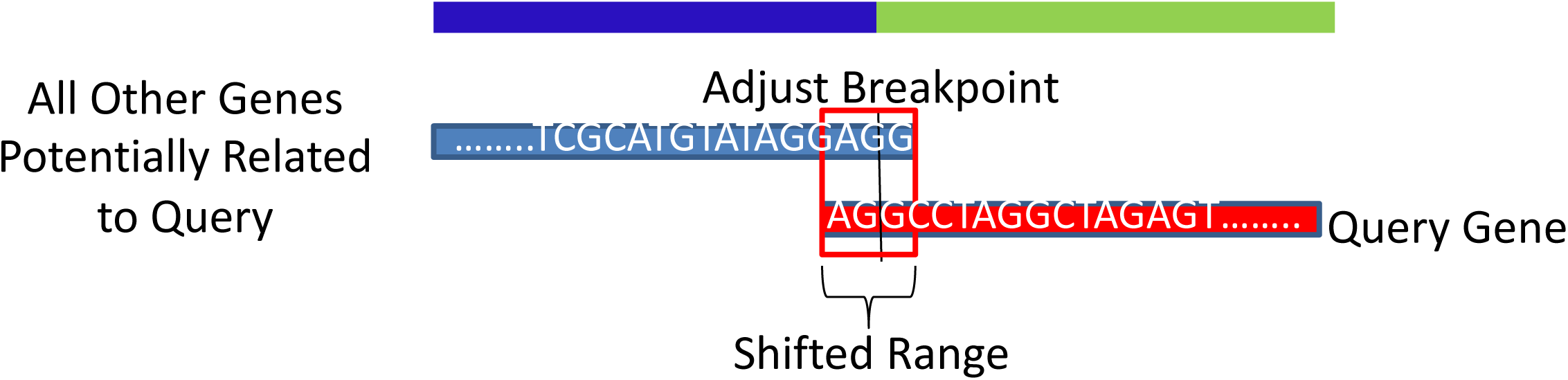
Breakpoint adjustment of the consensus sequence. When the consensus sequence is locally aligned to the query gene (red) and all other potential partners detected (blue), the two alignments can have a small overlap region (e.g. AGG) around the fusion breakpoint. Because the sequence of this region is the same on the two alignments, no matter where the fusion breakpoint is in this region (e.g. A-GG or AG-G), the fusion product is not changed. As a result, the size of the region is called shifted range and the breakpoint is adjusted to the middle of this region as the representative of the fusion event.

After adjusting fusion boundaries of fusion events, spanning reads (from paired-end processing category) were assigned to related fusion events. If a fusion event had *M* multiple-alignments, but only *M’* out of *M* had corresponding spanning reads, the multiple-alignment number was adjusted to *M’* instead of *M* (Figure 4). The rest of spanning reads, which were not assigned to any fusion events with splitting reads, were self-grouped into spanning-only fusion events by their alignment locations.

To reduce false positives, all the reported fusions were filtered by user defined parameters, including the minimum requirement of splitting, spanning, and total supporting reads.

To prioritize important and reliable fusions, seven features were used to rank and reorder the results:

1. Number of splitting reads.
2. Number of spanning read-pairs.
3. Total number of supporting reads (sum of splitting and spanning reads).
4. Degree of dispersion of splitting reads. In theory, a sample without PCR artifacts should have uniformly distributed splitting reads around the breakpoint (Figure S1). As a result, p-value from Kolmogorov-Smirnov (KS) test was used to evaluate how similar the distribution of splitting reads was to a uniform distribution.
5. Shifted range: the length of a ‘shifted region,’ in which shifting the breakpoint location would not change the fusion sequence. From literature, we have found that lots of validated fusions have ‘shifted regions.’^21^ As a result, we use ‘shifted range’ to evaluate the similarity around the breakpoint of the fusion gene pair. However, if the shifted range of a fusion was longer than 23bp, we considered it as a false positive caused by homology and filtered it out.
6. Sequence complexity around the breakpoint within 30 bp, measured by the dinucleotide entropy *E*: 

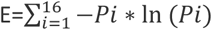 There were 4^2^=16 possible dinucleotide combinations. *P*_*i*_ denoted the frequency with which the *i*^*th*^ combination occured in the 30bp range. In the extreme scenario that the sequence was composed of only two types of dinucleotide, e.g. AAAAAAAAAAAAAAATATATATATATATAT, the entropy was (2×-0.5×ln(0.5)=0.69. Therefore, fusions with entropy ≤0.7 were considered as false positive and filtered.
7. Multiple-alignments number. Fusions with a single-alignment location were deemed most reliable. Conversely, the more alignments a fusion had, the less reliable it was.

Each fusion was assigned a score *S*_*rank*_ equal to the sum of the ranks of the features just described, S_rank_ = R_split_+ R_span_×2+ R_total_+ R_dispersion_+ R_shifted_range_+ R_complexity_+ R_multiple_alignment_, with R_(X)_ denoting the rank of the X feature (see supplement).

### RNA-Seq based fusion simulator – “SimFuse”^22^

SimFuse is an RNA-Seq based fusion simulator to evaluate and compare the performance of fusion-detection algorithms. With user-defined parameters, SimFuse randomly generates corresponding sample size with different numbers of fusion supporting reads. Using SimFuse simulated data, users can precisely estimate fusion detection recall and precision rates as a function of the numbers or types of supporting reads.

### In silico hypothesis testing tool for “broken” events – “PREF” ^23^

PREF is a novel *in silico* hypothesis testing tool for “broken” events (fusion, splicing and deletion) by generating putative references from hypothesized breaking locations.

## Datasets

### Simulated datasets

In this study, two simulated datasets were used. One was newly generated by a simulator called SimFuse, and the other one was from FusionMap^24^ as external validations.

In the SimFuse simulation, ENCODE MCF-7 cell line data (SRR521521) was used as the background. This cell line has a read length of 76 bp and mean fragment size of 192 bp. Thus, the theoretical ratio of splitting reads to spanning reads is 19:5, ideal for generating both splitting and spanning reads at the same time. In this simulated dataset, we generated 100 independent simulations, each of which had 10 different supporting read groups with 100 fusion events in each group (Table S1)^22^. The minimum-alignment length was set to 30 bp to ensure all the splitting reads were detectable by most algorithms.

The simulated dataset from FusionMap has read length of 75 bp and mean fragment size of 158 bp. It has 57209 paired-end reads and contains 50 simulated fusion events.

To obtain a fair comparison of results from different fusion-detection algorithms, the following criteria were applied.

1. Only the filtered results were compared (both deFuse and QueryFuse have filtered and unfiltered results).
2. Only chromosome, fusion boundaries, and fusion direction were used as “matching” criteria. Gene names and gene IDs were not used because there could be more than one gene at the same location.
3. Fusion direction also needed to be considered. For example, in any gene_1_-gene_2_ fusion, there could be only two distinct fusion events associated with the same fusion boundaries: gene_1_ on the 5’ end with gene_2_ on the 3’ end; and, vice versa, gene_1_ on the 3’ end with gene_2_ on the 5’ end.
4. To qualify as a true positive fusion, the fusion’s detected boundaries must be within the 10 bp range of its simulated boundaries.

Precision, rather than specificity, is used alongside recall (or sensitivity) to assess the accuracy of a fusion-detection algorithm. Precision is more informative than specificity, since in fusion detection the number of true negative (TN) is always much larger (∼20K genes in the human genome) than the number of true positive (TP). As a result, unless the number of false positive (i.e., type I errors) is extremely large, the specificity will always be close to 1, thus, will not adequately capture the difference among competing detection algorithms.

In this paper, QueryFuse v1.0, TopHatFusion v2.0.4 and deFuse v0.6.1 were used for comparisons.

### Cell line datasets

In addition to our internal data, we used 3 paired-end RNA-Seq cell line datasets with experimentally validated fusion as external test sets to estimate QF’s reliability in real data (Table S2).

1. Breast cancer cell line datasets^25^. BT-474, SK-BR-3 and MCF7 cell lines were used. The read length of this dataset is 50 bp, which is not optimal for QueryFuse. Datasets with longer read length (76 bp or 101 bp) for these 3 cell lines were found from DNAnexus Archive and used (Table S2).
2. ALK fusion cell line datasets^26^. H2228 cell line is a model of non-small cell lung cancer (NSCLC) with ALK-EML4 and ALK-PTPN3 fusion. DHL Cell line is a model of anaplastic large cell lymphoma (ALCL) with ALK-NPM1 fusion.
3. Bladder cancer cell line datasets. In SOAPFuse manuscript^27^, two bladder cancer cell lines were used to detect fusions and 15 fusion transcripts were experimentally confirmed.

Genes in the validated fusions were used as query genes for QueryFuse.

### Primary tumor specimens in lymphoma patients

In accordance with local IRB protocols, 16 fresh-frozen tumor samples from patients diagnosed with primary central nervous system lymphomas (10 patients) and primary testicular lymphomas (6 patients) were collected from 4 institutions (Dana-Farber Cancer Institute, Brigham and Women’s Hospital, Massachusetts General Hospital, all Boston, MA, USA; University of Freiburg, Germany^28^. High molecular weight genomic DNA was extracted from all frozen tumors. Matched total RNA was extracted from these 16 samples. We processed Whole Exon Sequencing (WES) on DNA and RNA-Seq on the matched RNA. RNA-Seq was run on Illumina® HiSeq 2000 to generate paired-end 100-bp reads with a median fragment size of 150bp.

### Implementation, availability and data resources

QueryFuse was implemented in shell script, python, perl, and R. A run with one query (60 kb) on 52 million paired-end reads was completed in approximately 215 min (80 min preprocessing and 135 min querying) using one core in an eight-core 2.6GHz Intel Xeon E5-2670 processor with 64GB memory. The preprocessing time and the querying time were dependent on read number, and the querying time was also dependent on the query gene length. QueryFuse was able to run different parallel queries with multiple processors.

The input reference genome (hg19) was downloaded from TopHat website. Gene annotation file of hg19 was downloaded from biomart (http://useast.ensembl.org/biomart/) with focus on protein-coding as gene type.

To rescue potential gene fusions with partners partially falling outside the gene annotation region, an expanded annotation file was generated from the regular gene annotation file. In this expanded annotation file, each gene was extended by 5,000 bp on both upstream and downstream.

QueryFuse is an open source package. All scripts and user manual are available in github (https://github.com/yuxiangtan/QueryFuse).

## RESULTS

### QueryFuse had the highest recall and precision rates in simulated datasets

To estimate both recall and precision rates^16^, we used SimFuse datasets (see datasets in Method Section) to compare performances of QueryFuse, TopHatFusion, and deFuse (Figure 6 and Table S3). QueryFuse had the highest recall and precision rates across all combinations of supporting reads. The highest recall rates achieved by QueryFuse, TopHatFusion, and deFuse were 90%, 84%, and 85%, respectively. Precision rates for QueryFuse and deFuse peaked at 99% and 95%, respectively. Of notice, TopHatFusion’s precision rate showed a slight decrease as the number of supporting read increased. This might be due to the fact that, for TopHatFusion the risk of detecting multiple-alignments increased as the number of supporting reads increased. Additionally, QueryFuse and TopHatFusion could detect fusions with as low as 4 supporting reads (3 splitting and 1 spanning). On the other hand, deFuse needed around 20 splitting and 5 spanning supporting reads to achieve a reasonable detection performance (75% recall and 95% precision rates).

**Figure 6.**
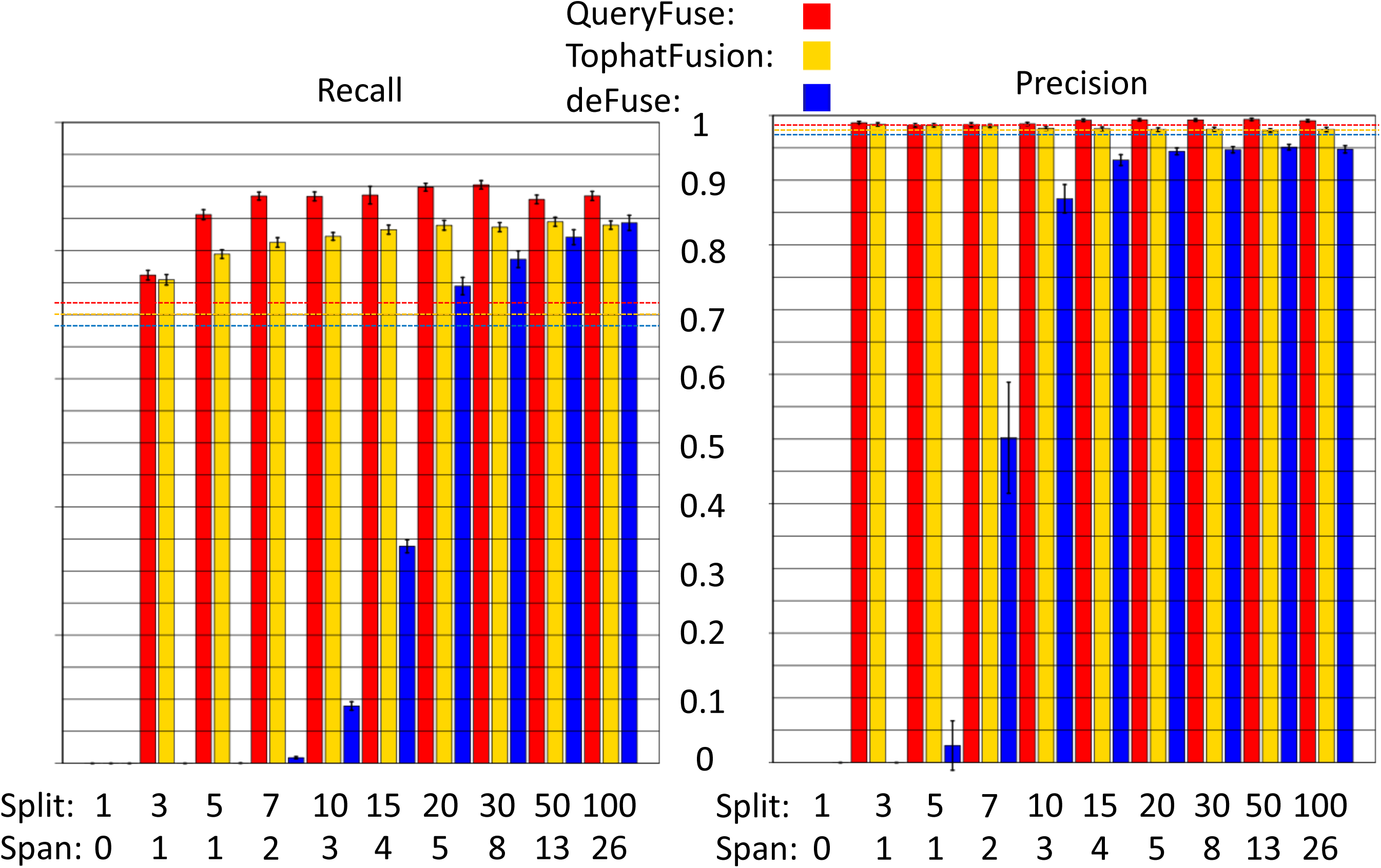
Barplot of recall and precision for QueryFuse, TopHatFusion and deFuse. Red bars indicate QueryFuse results; blue bars indicate deFuse results: and yellow bars indicate TopHatFusion results. The x-axis is indexed by the supporting read groups with each group corresponding to the indicated splitting-to-spanning read ratio. The y-axis reports recall and precision rates. The upper and lower whiskers represent the upper and lower limits of the confidence interval at 95% confidence level. The dashed line shows estimates from the FusionMap dataset. The red dashed line shows estimates for QueryFuse; the yellow line shows estimates for TopHatFusion; and the blue line shows estimates for deFuse.

For the simulation dataset from FusionMap, QueryFuse had the highest recall and precision rates (36/50=72%) and (36/37=97.3%), respectively. TopHatFusion had the second highest recall and precision rates (35/50=70%) and (35/36=97.2%) respectively. deFuse’s recall and precision rates were (34/50=68%) and (34/35=97.1%) respectively.

### QueryFuse detected most of the experimental validated fusions in cell line datasets

For breast cancer datasets, the numbers of experimental validated fusions in cell lines were 10 in BT-474, 7 in SK-BR-3, and 3 in MCF7 ^25^ (Table S2). However, in the DNAnexus Archive datasets with longer read lengths, many validated fusions were missed by all three algorithms (6 in BT-474, 3 in SK-BR-3, and 0 in MCF7) (Table S2). This showed potential batch effects from a variety of cells in the cell lines. QueryFuse worked as well as deFuse in detecting fusions (3 in BT-474, 3 in SK-BR-3, and 3 in MCF7), while TopHatFusion detected one more fusion in SK-BR-3 than the others. CCDC85C-SETD3 fusion was also detected in SK-BR-3 by QueryFuse, but supported only by spanning reads. Since CCDC85C-SETD3 fusion had a low supporting read number, it is possible that no detectable splitting reads of this fusion exist. Because the theoretical splitting:spanning read ratio of the 50-bp-long datasets is 1:1, it is less likely to detect splitting reads than the DNAnexus Archive datasets. Thus, fusion events supported by only spanning reads in QueryFuse should also be considered as positive events. QueryFuse performed equally well as deFuse (7 in BT-474, 4 in SK-BR-3, and 3 in MCF7), while TopHatFusion (5 in BT-474, 4 in SK-BR-3, and 2 in MCF7) missed 3 events (Table S2). The validated fusions missed by all three algorithms (2 in BT-474, 2 in SK-BR-3, and 0 in MCF7) had low supporting read numbers, which made them undetectable.

In ALK dataset, all five H2228 samples have ALK-EML4 and ALK-PTPN3 fusions and three DHL samples have ALK-NPM1 fusion. deFuse detected all these fusions, while QueryFuse missed one fusion event and TopHatFusion missed three fusion events.

In the bladder cancer cell line dataset, deFuse, TopHatFusion, and QueryFuse performed equally well (3 out 8 in SRR497884 and 3 out 7 in SRR497885).

In summary, although it is challenging to know all the true positives and true negatives of fusions in real datasets, we used the number of detected fusions from the validated fusions to evaluate the sensitivity of deFuse, TopHatFusion, and QueryFuse. For all the candidate fusion events, deFuse and QueryFuse detected the same number of events (42 out of 68), while TopHatFusion detected 37.

### QueryFuse found all isoforms of a hypothesized fusion in a clinical dataset

Because of partial copy number gain and upregulated expression on PDCD1LG2 in the clinical sample TL-03, we hypothesized that PDCD1LG2 was involved in a fusion event with replacement of regulation elements^28^. Using QueryFuse to query on PDCD1LG2, we found the hypothesized fusion with three RNA isoforms in this sample (Table 2)^28^. All the isoforms supported the same DNA fusion event in which upstream regulation elements of PDCD1LG2 were replaced by elements from TBL1XR1. Additionally, we ran two global fusion-detection algorithms (deFuse and TopHatFusion) to validate this finding and compared it with QueryFuse. deFuse found two out of three isoforms of this fusion, while TopHatFusion totally missed all three isoforms (Table 2). This fusion was experimentally validated by 5’RACE PCR^28^. The PCR result showed the isoform was discovered by QueryFuse, but not by deFuse (bold in Table 2). These three isoforms were *in silico* validated by PREF^23^.

**Table 2.** Hypothesized *PDCD1LG2* related fusion isoforms found in a lymphoma patient

| Fusion Gene (Partner) | Breakpoint (Partner) | Fusion Gene (Query) | Breakpoint (Query) | Supporting Read # | Reported from deFuse | Reported from TopHatFusion |
| --- | --- | --- | --- | --- | --- | --- |
| *TBL1XR1* | 176914912 | *PDCD1LG2* | 5522529 | 56 | Yes | No |
| ***TBL1XR1*** | **176878662** | ***PDCD1LG2*** | **5522529** | **29** | **No** | **No** |
| *TBL1XR1* | 176914129 | *PDCD1LG2* | 5522531 | 10 | Yes | No |

From the WES data, we detected seven fusions which were in the in-frame regions of genes^28^. As a result, we used deFuse, TopHatFusion, and QueryFuse on matched RNA-Seq data to check the presence of these fusions (Table S4). QueryFuse and deFuse detected the same amount of events from these fusion events, while TopHatFusion detected none.

## DISCUSSION

### QueryFuse was sensitive in both simulated datasets and real datasets

Overall, among these three methods in both simulated and real datasets, QueryFuse had the highest number of detecting TPs. In the simulated dataset from SimFuse, QueryFuse had the best recall and precision rates for all combinations of supporting reads. In the simulated dataset from FusionMap, QueryFuse had the best recall and precision rates, as well. Although the differences in performance of cell line datasets among the three methods were not as significant as in simulated datasets, QueryFuse’s performance was one of the foremost. Furthermore, in the clinical dataset, only QueryFuse could detect an extra RNA isoform from the fusion event, which was experimentally validated. This showed that QueryFuse was potentially more sensitive in detecting RNA fusion isoforms than global detection methods. However, this hypothesis is difficult to validate and further research is necessary.

### QueryFuse had proven to be the most appropriate algorithm for each study of fusion hypothesis on a gene list

QueryFuse is a novel algorithm focusing on gene-specific fusion detection from RNA-Seq data. Its primary strength is to test fusion hypotheses on a list of genes reliably and efficiently. With aligned RNA-Seq data, users can check their fusion hypotheses even before spending time on experimental validations. By integrating copy number changes with gene expression changes, biologists can raise hypotheses of fusion genes^28^. With DNA-Seq and matched RNA-Seq data, researchers can check whether a fusion detected in DNA-Seq is expressed in RNA-Seq^28^. Therefore, gene-specific fusion detection is more appropriate than global fusion detection and QueryFuse is designed to meet this requirement. Additionally, it is inefficient for biologists to peruse a long list of fusions from the results of a global fusion detection tool, because most of the fusions are unrelated to their studies. Therefore, by using QueryFuse, researchers can fully focus on relevant genes and prioritize the most reliable fusions through its ranked results.

### QueryFuse performed more rapidly and efficiently than TopHatFusion and deFuse when the query gene list was less than 16 genes

In the 52 million paired-end reads example, to run one query in a single processor, QueryFuse took 215 min (80 min preprocessing time + 135 min querying time); in comparison, deFuse took 1398 min and TopHatFusion took 3075 min for a genome-wide detection. This showed the computational advantage of using the QueryFuse algorithm. It was noted that running at parallel mode saved time for deFuse and TopHatFusion. For example, running in 16 parallel processors, the same sample took 150 min for deFuse and 516 min for TopHatFusion. Although QueryFuse could not run parallel for a single query, for *X* queries, QueryFuse could run in *X* processors with similar processing times. For a test with 16 random queries in this example in 16 processors, QueryFuse took 170 min (80 min preprocessing time + 90 (±102) min querying time). As a result, with the same computational resource, for a list of 16 query genes, QueryFuse ran at similar times as deFuse. Additionally, because the querying time was almost linearly related to the length of the query gene (Figure S2), QueryFuse performed efficiently on short query genes.

### Future Improvements

QueryFuse, as a gene-specific algorithm, made use of the known query gene to narrow down fusion targets. Because it targeted and used extra information from splitting reads, QueryFuse was more efficient than existing methods, which was validated in a single core run. Additionally, we showed that QueryFuse was faster than other global fusion detection methods when the query gene list was less than 16 genes. However, QueryFuse’s implementation was not computationally efficient enough on parallelization and requires room for improvement.

Meanwhile, the sensitivity and specificity of BLAT dropped significantly when the reads were short (less than 30 bp). As a result, replacing BLAT with better local aligners should improve QueryFuse’s performance (recall and precision), which requires further testing.

At this moment, QueryFuse’s input must be the aligned BAM file from fusion-detection free RNA-Seq aligners, such as TopHat, SOAP, etc. However, it was noted that some new aligners, such as STAR, have integrated fusion-detection as default parameters in the regular read alignment step. Because fusion supporting reads from these integrated aligners are annotated differently, in the future, a complementary option will be included in QueryFuse to choose type of input.

With the development of sequencing technology, long single-end sequencing might become popular. Although QueryFuse can presently only work on paired-end sequencing data, it should theoretically work for single-end sequencing as well, which requires additional improvements.

## ACKNOWLEDGEMENT

The authors would like to thank Stefano Monti, Bjoern Chapuy, Avi Spira, Evan Johnson, Luis Carvalho, Liye Zhang, Yann Tambouret, and Daniel Gusenleitner for data and feedbacks for improvement. The authors would also like to thank Charron Cote, Jake Kantrowitz and Yi Li, as well as the anonymous reviews for their valuable comments and suggestions to improve the manuscript.

## FIGURE LEGEND

**Fig S1.**
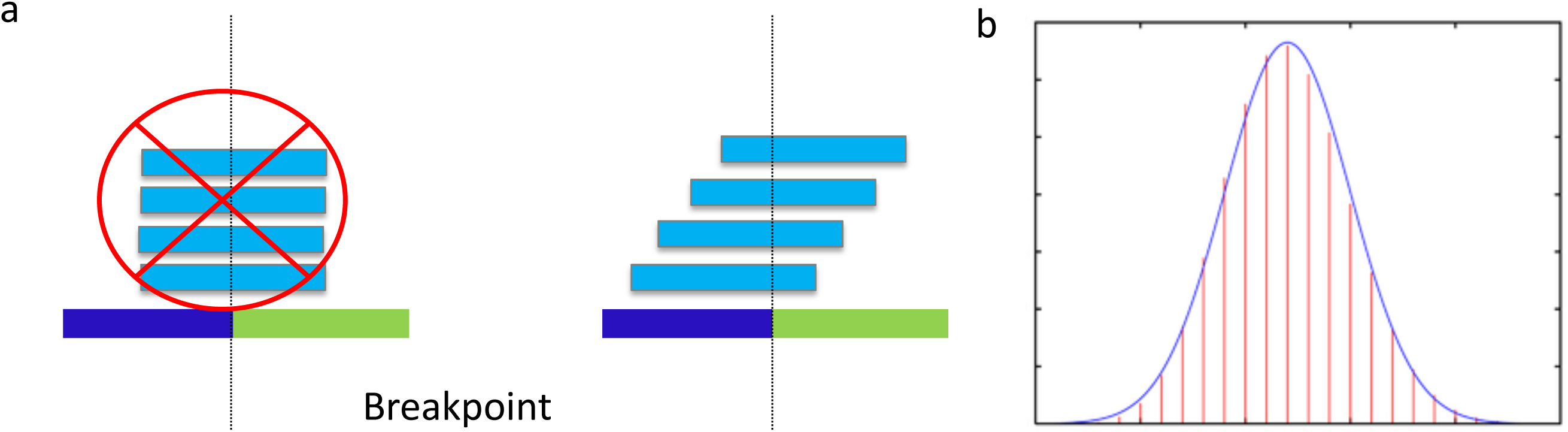
Dispersion of splitting reads.

**Fig S2.**
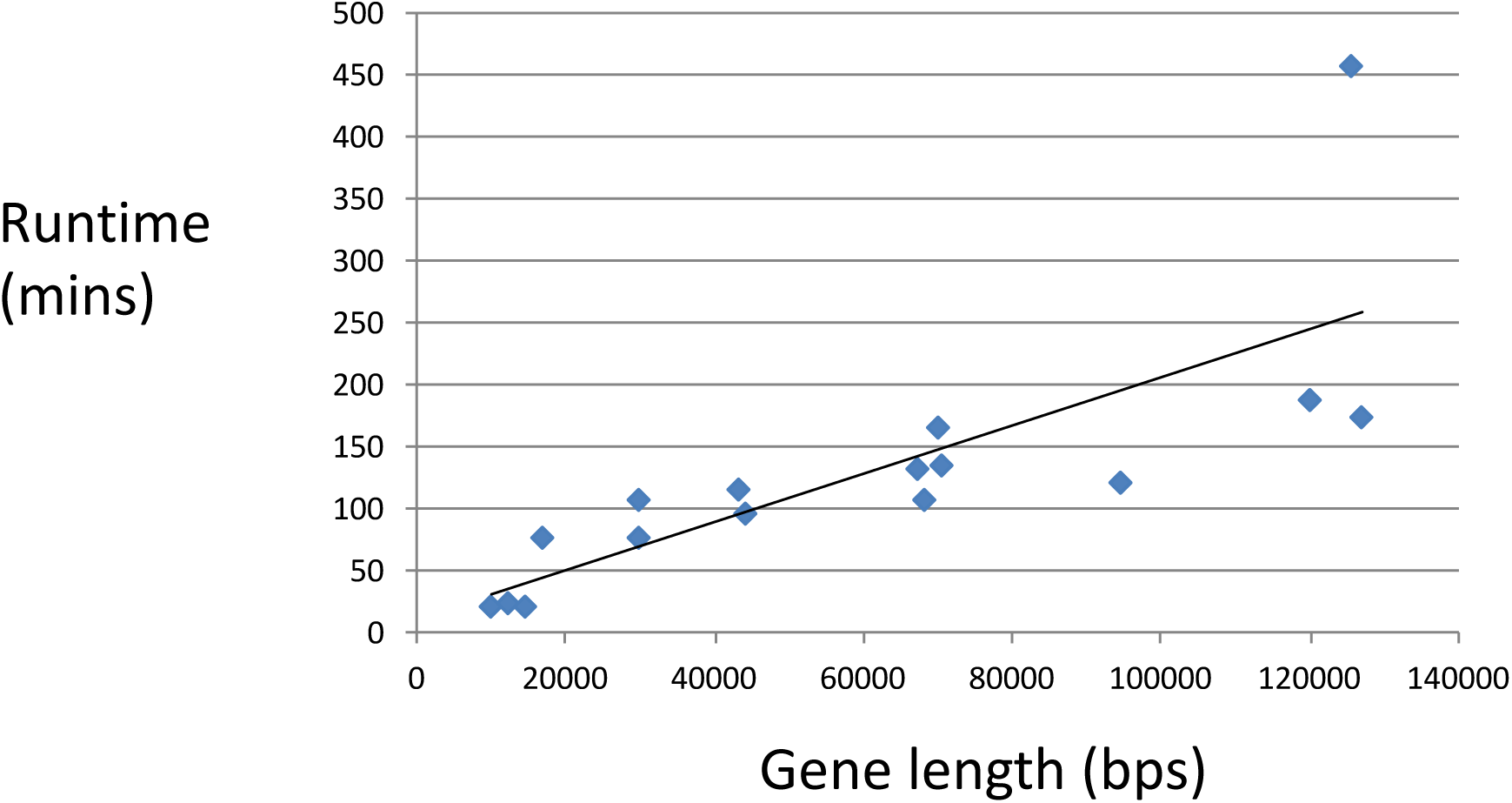
Run time and gene length relation

**Fig S3.**
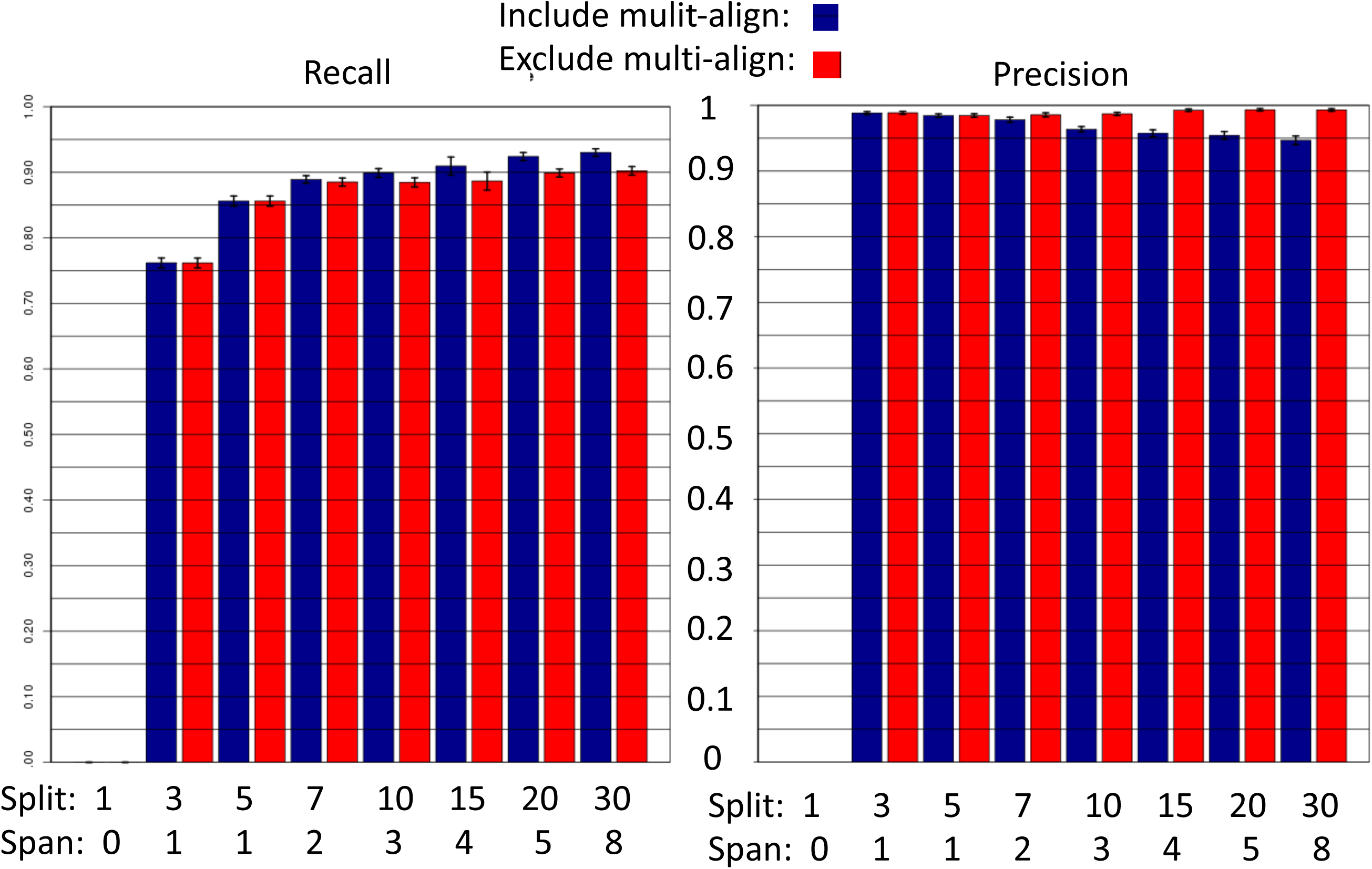
Recall and precision change in QueryFuse with and without allowing multiple-alignment.

