## Supplemental file1 for "Queryfuse is a sensitive algorithm for detection of gene-specific fusions"

### Supplementary Computational Methods

#### Result summarization

##### Spanning-only fusion events

If a spanning-only fusion event has location *A* and location *B* as the fusion boundaries, only the spanning read pairs, in which one end is within the range of *A*±read-length×2 and another end is within the range of *B*±read-length×2, are grouped as supporting reads for this event.

##### Result ranking and scoring

Each fusion is assigned a score *S_rank_* equal to the sum of the ranks of seven features.

S_rank_ = R_split_+ R_span_*2+ R_total_+ R_dispersion_+ R_shifted_range_+ R_complexity_+ R_multiple_alignment_

The rank of spanning read number (R_span_) has weight of 2 because generally R_span_ is twice important as the other features. This is because in regular library preparation procedure, the insert size is negative, which make fusions with spanning reads a strong indicator of true positives.

### Supplementary Results and Discussion

#### Comparison of recall and precision rates of QueryFuse between the setting allowing multiple-alignment and not

In figure S3, we compared QueryFuse results with and without multiple-alignment events from the SimFuse data. If fusions with multiple-alignment are included in the final report, a fusion with multiple-alignment (generally they are from the homologous region) will have multiple entries, which show different breakpoint locations, in the result. However, in the simulation process, although the homologous stages of fusion genes are not checked, all fusions are assumed to be uniquely aligned. (This assumption is not always correct and requires further improvement to fix.) As a result, these additional multiple-alignment results were falsely considered false positive and decreased the precision rate. Additionally, the chance to detect fusions with multiple-alignment is lower in the fusions with small supporting read number than the fusions with large supporting read number, because fusions with multiple-alignment requires more supporting evidences than the fusions with unique-alignment. This led to the strange situation that the precision rates decreased as the supporting read number increased. To correct this estimation, we used QueryFuse results in which fusions with multiple-alignment were excluded. In this case, although the recall rate was slightly lower than the case in which fusions with multiple-alignment were included, the precision rate increased as the supporting read number increased.

In fact, from QueryFuse’s final output, users can recognize the fusions with multiple-alignment by checking the group number, which won’t lead to the conclusion of false positive. As a result, the default mode of QueryFuse allows fusions with multiple-alignment in the report because users can benefit from the additional information. On the other hand, in deFuse, only the most likely set of fusion events are reported for ambiguous alignments. However, for homologous regions, in most cases, all the events are equally reliable because their sequences are just the same. This makes deFuse’s result misleading on these cases. In TophatFusion, homologous events are reported multiple times as QueryFuse, but they are not marked as multiple-alignment, which is misleading as well. However, in this paper, to simplify the comparison among three methods, all fusions with multiple-alignment were excluded from the QueryFuse’s output.

### Supplementary Table Legend

Supplementary table 1: Simulation matrix I used to generate the SimFuse example dataset.

Supplementary table 2: Fusion detection result on cell lines

Supplementary table 3: Recall and precision statistics among QueryFuse, deFuse and TophatFusion

Supplementary table 4: Validation of detected fusion from WES data by using match RNA-Seq data with QueryFuse, deFuse and TophatFusion.

### Supplementary Figure Legend

Supplementary figure 1: Dispersion of splitting reads. a) Dash line is the fusion breakpoint. On the left, all the supporting reads are piled up at the same location, which is the evidence of PCR artifacts and make the fusion less reliable. On the right, all the supporting reads are well spread, which indicates no PCR artifacts. b) Theoretical distribution of coverages on each base, which should be a normal distribution.

Supplementary figure 2: Liner relation between gene length and query time. X axis is the length of query genes and Y axis is the runtime on each query (query time).

Supplementary figure 3: Barplot of recall and precision rates of QueryFuse with and without multiple-alignment results. Blue bars stand for QueryFuse results which includes fusions marked with multiple-alignment. Red bars stand for QueryFuse results which excludes fusions marked with multiple-alignment. X axis is supporting read number groups. Each group is composed by a splitting read number and spanning read number. Y axis is rates.
